## Supporting Information for "Bayesian active learning for optimization and uncertainty quantification in protein docking"

Yue Cao and Yang Shen
Department of Electrical and Computer Engineering
Texas A&M University
College Station, TX 77843-3128, US

#### 1 Theory

We first theoretically and empirically compare our Bayesian active learning (BAL) to [Ortega et al., 2012] (NCPD for Nonparametric Conjugate Prior Distribution) that also models the posterior of the global optimum directly. During the comparison, we establish BAL's advantages in theory, namely (1) the annealing schedule balancing exploration and exploitation is aware of the global uncertainty and dependent on dimensionality of the search space; and (2) the Kriging regressor is consistent and unbiased. We also establish BAL's advantage in practice through empirical comparison over test functions.

##### 1.1 Unlike NCPD, BAL's annealing schedule is global uncertainty-aware and dimension-dependent

In Nonparametric Conjugate Prior Distribution, the temperature constant $\rho$ was estimated proportional to the effective number of data points:

$$\rho_{\mathrm{NCPD}} = \rho_0(\xi + n \cdot \frac{\sum_i K(x_i, x_i)}{\sum_i \sum_j K(x_i, x_j)})$$

where $\rho_0$ is the initial value for $\rho$, and $\xi$ is the effective number of data points in the prior distribution and $n$ is the number of data points, which is penalized by $\frac{\sum_i K(x_i,x_i)}{\sum_i \sum_j K(x_i,x_j)}$ to become the effective number of sample points. The rational is that the effective number of samples somehow measures the uncertainty in the system.

However, as our problem has a constraint for the search space, only considering the pairwise distance between samples is obviously insufficient. The location of samples in the search space also contributes to the global uncertainty in the system. Considering the two cases in Fig. S1a and Fig. S1b. Here we have three data points which have the exact same pairwise distance between each other. The figure shows the standard deviation within the square search space. Obviously, the second situation has more uncertainties than the first one because the points in the second figure are closer to the boundary, which makes the large region of the right bottom untouched and high variance. But the $\rho$ used in NPCD remains the same for both cases.
In contrast, the $\rho$ in our BAL is defined as

$$\rho_{\mathrm{BAL}} = \rho_0 \cdot \exp((h_p^{(t-1)})^{-1} n_t^{\frac{1}{d}})$$

Here we use $h_p^{(t-1)}$, the continuous entropy for the latest posterior distribution $p$, which is a global measure of uncertainty. In other words, we consider not only the internal structure between the samples but also the location of samples within the search space. Obviously our $\rho$ for the case in Fig. S1a is bigger than that for Fig. S1b, which makes much more sense.
Moreover, a good temperature constant should be generalizable for different dimensions. However, NCPD is bad-generalized for different dimensions. For instance, we remain the same pairwise distance between any two samples as shown Fig. S1a and then we extends the dimension from 2 to 8. It could be seen

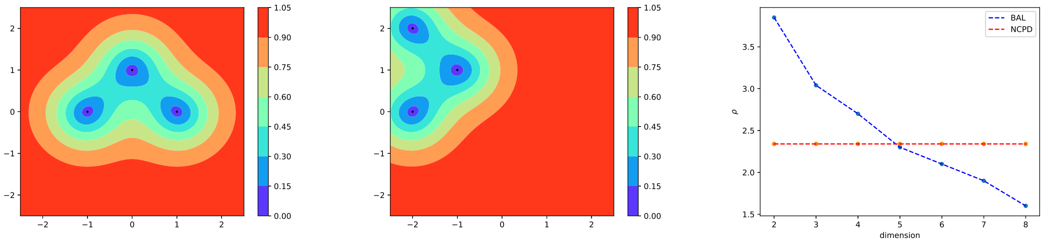


(a) (b) (c)

Figure S1: (a): The contour plot for the standard deviation within [-2.5,2.5]×[-2.5,2.5] with three data points at [-1,0], [1,0], [0,1]. (b) The contour plot for the standard deviation within [-2.5,2.5]×[-2.5,2.5] with three data points at [-2,0], [-2,2], [-1,1]. (c) The $\rho$ keeps unchanged for NCPD(red line) as dimension goes higher, while it decreases quickly in BAL (blue line).

in Fig. S1c. The $\rho$ in NCPD remains the same while in BAL it decreases rapidly as the dimension goes larger. For the same set of data, as the dimension goes higher, the uncertainty of the system must decrease, which means $\rho$ must decrease at the same time. Therefore, compared to NCPD our BAL matches the rationale and could be generalizable for various dimensions.

Lastly, we found that the $\rho$ in NCPD decreases as the number of samples increases in some situations. This is totally controversial to our rational for $\rho$. As we are getting more samples, our knowledge about the system is increasing (not dropping the old samples). In adaptive simulated annealing, this means our system is getting cooler and cooler as the annealing procedure goes forward. Therefore, there is a monotonous-positive relationship between $\rho$ and the number of samples. However, considering the simple example below, we have two data points $\boldsymbol{x_1} = [1, 0]$ and $\boldsymbol{x_2} = [-1, 0]$. We considered a kernel that $K(\boldsymbol{x_1}, \boldsymbol{x_2}) \approx 0$ and $K(\boldsymbol{x_i}, \boldsymbol{x_i}) = 1$(e.g. RBF kernel with bandwith $l << 1$). The effective number of location for the sample is

$$t_2 = 2 \cdot \frac{\sum_i K(x_i, x_i)}{\sum_i \sum_j K(x_i, x_j)} = 2 \cdot \frac{2}{2} = 2$$

Then we add the third sample $\boldsymbol{x_3} = [-1, 0 + \epsilon]$, where $\epsilon$ is a tiny positive number, so that $K(\boldsymbol{x_1}, \boldsymbol{x_3}) \approx 0$ and $K(\boldsymbol{x_1}, \boldsymbol{x_2}) \approx 1$. Then the effective number of location will become:

$$t_3 = 3 \cdot \frac{\sum_i K(x_i, x_i)}{\sum_i \sum_j K(x_i, x_j)} = 3 \cdot \frac{3}{5} = 1.8$$

We have $t_3 < t_2$! That means when collecting a new sample $\boldsymbol{x_3}$, the system's uncertainty is becoming larger. Although the new sample is located very closed to the old one, it is obviously controversial to our understanding and rationale for $\rho$. The situation for $\rho$ decreasing frequently happens when n becomes large, resulting from the new samples having a large chance to be closed to the old ones. By contrast, in BAL, our $\rho$ is in positive relation to $n$ and the negative relation to $H$, while $n$ is getting larger and $H$ is getting smaller over iterations. Therefore, our $\rho$ strictly increased when the system is getting more samples.

#### 1.2 Unlike NCPD, BAL's Kriging regressor is unbiased

In this section we prove the regressor used in NCPD is biased. Then we follow [Matheron, 1963] and briefly derive our Kriging results.

##### 1.2.1 The regressor in NCPD is biased

We first consider the one-dimensional case, and then extend to multi-dimensional cases. Suppose we have two groups of random variables $X_i, Y_i$ which are *i.i.d.* with joint pdf (probability distribution function) $p(x, y)$. The mean function $f(x) = E[Y|X = x]$ is the true function that we want to predict. Suppose the marginal pdf of $X$ has the form

$$p(x) \propto e^{\frac{f(x)}{\beta}}$$

The estimator in NCPD for $f(x)$ is

$$\widehat{f}(x) = \frac{\sum_{i=1}^{n} K(\frac{x-x_i}{h})y_i + k_0(x) * y_0(x)}{\sum_{i=1}^{n} K(\frac{x-x_i}{h}) + k_0(x)}$$

For a given $x_i$, the random variable $y_i$ can be written as

$$y_i = f(x_i) + \epsilon_i$$

where $\epsilon_i$ is a zero-mean noise. In our case, it could be regarded as the system error and the difference between our supposed energy function and the true energy function.

Therefore, we have

$$\sum_{i=1}^{n} K(\frac{x-x_i}{h})y_i = \sum_{i=1}^{n} K(\frac{x-x_i}{h})f(x_i) + \sum_{i=1}^{n} K(\frac{x-x_i}{h})\epsilon_i$$

add prior function to both sides, and divide them by $\sum_{i=1}^{n} K(\frac{x-x_i}{h}) + k_0(x)$, we reach

$$\frac{\sum_{i=1}^{n} K(\frac{x-x_i}{h})y_i + k_0(x) * y_0(x)}{\sum_{i=1}^{n} K(\frac{x-x_i}{h}) + k_0(x)} = \frac{\sum_{i=1}^{n} K(\frac{x-x_i}{h})f(x_i) + \sum_{i=1}^{n} K(\frac{x-x_i}{h})\epsilon_i + k_0(x) * y_0(x)}{\sum_{i=1}^{n} K(\frac{x-x_i}{h}) + k_0(x)}$$

Note that the left side is just $\widehat{f}(x)$. So we have

$$\widehat{f}(x) - f(x) = \frac{\sum_{i=1}^{n} K(\frac{x-x_i}{h})f(x_i) + \sum_{i=1}^{n} K(\frac{x-x_i}{h})\epsilon_i + k_0(x) * y_0(x)}{\sum_{i=1}^{n} K(\frac{x-x_i}{h}) + k_0(x)} - f(x)$$

Multiply both the nominator and the denominator by $\frac{1}{nh}$, we have

$$\widehat{f}(x) - f(x) = \frac{\frac{1}{nh}\sum_{i=1}^{n} K(\frac{x-x_i}{h})f(x_i) + \frac{1}{nh}\sum_{i=1}^{n} K(\frac{x-x_i}{h})\epsilon_i + \frac{1}{nh}k_0(x) * y_0(x)}{\frac{1}{nh}\sum_{i=1}^{n} K(\frac{x-x_i}{h}) + \frac{1}{nh}k_0(x)} - f(x)$$

Note the the first term in the denominator is the kernel density estimator for $p(x)$

$$\widehat{p}(x) = \frac{1}{nh}\sum_{i=1}^{n} K(\frac{x-x_i}{h})$$

We can rewrite the whole equation

$$\widehat{f}(x) - f(x) = \frac{\frac{1}{nh}\sum_{i=1}^{n} K(\frac{x-x_i}{h})(f(x_i) - f(x)) + \frac{1}{nh}\sum_{i=1}^{n} K(\frac{x-x_i}{h})\epsilon_i + \frac{1}{nh}k_0(x) * (y_0(x) - f(x))}{\widehat{p}(x) + \frac{1}{nh}k_0(x)}$$

Define

$$m_1(x) = \frac{1}{nh}\sum_{i=1}^{n} K(\frac{x-x_i}{h})(f(x_i) - f(x))$$

$$m_2(x) = \frac{1}{nh}\sum_{i=1}^{n} K(\frac{x-x_i}{h})\epsilon_i$$

We can simplify the equation above as

$$\widehat{f}(x) - f(x) = \frac{m_1(x)}{\widehat{p}(x) + \frac{1}{nh}k_0(x)} + \frac{m_2(x)}{\widehat{p}(x) + \frac{1}{nh}k_0(x)} + \frac{1}{nh}\frac{k_0(x) * (y_0(x) - f(x))}{\widehat{p}(x) + \frac{1}{nh}k_0(x)}$$

By fixing $x$, we start to analyze each term in the right side of the equation.

(1) We start with the simplest term among the three, the second term. We first calculate the expectation and variance of $m_2(x)$.
Since we know $E[\epsilon_i|X_i] = 0$

Then

$$E[m_2(x)] = E_{x_i}[E_{\epsilon_i|x_i}[\frac{1}{nh}\sum_{i=1}^{n} K(\frac{x-x_i}{h})\epsilon_i]] = E_{x_i}[0] = 0$$

Because $(X_i, Y_i)$ are i.i.d, we have

$$Var[m_2(x)] = \frac{1}{n^2h^2}\sum_{i=1}^{n} Var[K(\frac{x-x_i}{h})\epsilon_i] = \frac{1}{nh^2}E[K^2(\frac{x-x_i}{h})\epsilon_i^2]$$

Define $\sigma^2(x) = E[\epsilon_i^2|X_i]$. We have

$$Var[m_2(x)] = \frac{1}{nh^2}E[K^2(\frac{x-x_i}{h})\sigma^2(x)] = \frac{1}{nh^2}\int K^2(\frac{x-x_i}{h})\sigma^2(x_i)p(x_i)dx_i$$

By setting $u = \frac{x_i-x}{h}$, we have

$$Var[m_2(x)] = \frac{1}{nh}\int K^2(u)\sigma^2(hu+x)p(hu+x)du$$

We use Taylor expansion for $\sigma^2(hu+x)$ and $p(hu+x)$ up to $o(h)$, and get

$$Var[m_2(x)] = \frac{1}{nh}\int K^2(u)\sigma^2(x)p(x)du + o(\frac{1}{n})$$

Define $\mu = \int K^2(u)du$. We obtain

$$Var[m_2(x)] = \frac{\mu\sigma^2(x)p(x)}{nh} + o(\frac{1}{n})$$

By applying Central Limit Theorem, we get the asymptotic result for $m_2(x)$

$$\lim_{\substack{h\to 0\\ nh\to\infty}} \sqrt{nh}m_2(x) \longrightarrow_d N(0, \mu\sigma^2(x)p(x))$$

(2) Second, we work on the first term and calculate the expectation and variance of $m_1(x)$. Because $X_i s$ are *i.i.d*, we have

$$E[m_1(x)] = \frac{1}{h}E[K(\frac{x-x_i}{h})(f(x_i)-f(x))] = \frac{1}{h}\int K(\frac{x-x_i}{h})(f(x_i)-f(x))p(x_i)dx_i$$

Let $u = \frac{x_i-x}{h}$,

$$E[m_1(x)] = \int K(u)(f(hu+x)-f(x))p(hu+x)du$$

Similar to the work above, we expand $(f(hu+x)-f(x))$ and $p(hu+x)$ up to $o(h^2)$, and obtain

$$\begin{aligned}E[m_1(x)] &= \int K(u)(huf^{'}(x) + \frac{h^2u^2}{2}f^{''}(x))(p(x)+hup^{'}(x))du + o(h^3)\\ &= hf^{'}(x)p(x)\int K(u)udu + h^2(f^{'}(x)p^{'}(x) + \frac{1}{2}f^{''}(x)p(x))\int K(u)u^2du + o(h^3)\end{aligned}$$

Let $\kappa_2 = \int K(u)u^2du$. Since $\int K(u)udu = 0$, we have

$$E[m_1(x)] = h^2\kappa_2(f^{'}(x)p^{'}(x) + \frac{1}{2}f^{''}(x)p(x)) + o(h^3)$$

The same method can obtain

$$Var[m_1(x)] = o(\frac{h^2}{nh})$$

Therefore, we have

$$\lim_{\substack{h\to 0\\ nh\to\infty}} \sqrt{nh}(m_1(x) - h^2\kappa_2(f^{'}(x)p^{'}(x) + \frac{1}{2}f^{''}(x)p(x))) \longrightarrow_p 0$$

The kernel density estimator $\widehat{p}(x)$ has the property

$$\lim_{\substack{h\to 0\\ nh\to\infty}} \widehat{p}(x) \longrightarrow_p p(x)$$

By using Slutskys theorem, and let $B(x) = f^{'}(x)p^{'}(x)p^{-1}(x) + \frac{1}{2}f^{''}(x)$, we get

$$\lim_{\substack{h\to 0\\ nh\to\infty}} \sqrt{nh}(\frac{m_1(x)}{\widehat{p}(x) + \frac{1}{nh}k_0(x)} + \frac{m_2(x)}{\widehat{p}(x) + \frac{1}{nh}k_0(x)} - h^2\kappa_2 B(x)) = \lim_{\substack{h\to 0\\ nh\to\infty}} \sqrt{nh}(\frac{m_1(x) + m_2(x)}{p(x)})$$

$$\longrightarrow_d N(0, \frac{\mu\sigma^2(x)}{p(x)})$$

(3) Third, we calculate the last term in $\widehat{f}(x) - f(x)$. Note that $\widehat{p}(x)$ is the only part including random variables of $\frac{1}{nh}\frac{k_0(x)*(y_0(x)-f(x))}{p(x)+\frac{1}{nh}k_0(x)}$. So we can easily conclude

$$\lim_{\substack{h\to 0\\ nh\to\infty}} \frac{k_0(x) * (y_0(x) - f(x))}{\widehat{p}(x) + \frac{1}{nh}k_0(x)} \longrightarrow_p \frac{k_0(x) * (y_0(x) - f(x))}{p(x)}$$

so that

$$\lim_{\substack{h\to 0\\ nh\to\infty}} \sqrt{nh}(\frac{1}{nh}\frac{k_0(x) * (y_0(x) - f(x))}{\widehat{p}(x) + \frac{1}{nh}k_0(x)}) \longrightarrow_p 0$$

Therefore, in summary, we obtain

$$\lim_{\substack{h\to 0\\ nh\to\infty}} \sqrt{nh}(\widehat{f}(x) - f(x) - h^2\kappa_2 B(x)) \longrightarrow_d N(0, \frac{\mu\sigma^2(x)}{p(x)})$$

It is easy to extend the result to multi-variable cases: For $d$ dimensions, we have

$$\lim_{\substack{H\to 0\\ n|H|\to\infty}} \sqrt{n|H|}(\widehat{f}(\mathbf{x}) - f(\mathbf{x}) - \kappa_2\sum_{i=1}^{i=d} h_i^2 B_i(\mathbf{x})) \longrightarrow_d N(0, \frac{\mu^d\sigma^2(\mathbf{x})}{p(\mathbf{x})})$$

where $H$ is the bandwidth matrix. In the Gaussian kernel, it is the covariance matrix.

In total, we have proved that the regressor in NCPD is biased and converges to the normal distribution with mean equal to $\kappa_2\sum_{i=1}^{i=d} h_i^2 B_i(\mathbf{x})$.

##### 1.2.2 Derivation of BAL's Kriging regressor

We briefly derive the Kriging regressor following [Chilès and Delfiner, 2012]. Let $F(\boldsymbol{x})$ be a random function with mean equal to $f(\boldsymbol{x})$, and $\{\boldsymbol{x}_i, y_i\}_{i=1}^n$ be our observed data with a noise $\epsilon$. We are trying to find an unbiased linear estimator $\hat{f}(\boldsymbol{x}) = \sum_i \lambda_i(\boldsymbol{x})y_i$ for $f(\boldsymbol{x})$ with the smallest variance:

$$\text{Minimize } Var[(\hat{f}(\boldsymbol{x}) - F(\boldsymbol{x}))^2]$$
$$\text{Subject to } E[\hat{f}(\boldsymbol{x})] = E[F(\boldsymbol{x})]$$

We define the covariance between $f(\boldsymbol{x}_1)$ and $f(\boldsymbol{x}_2)$ as $cov(\boldsymbol{x}_1, \boldsymbol{x}_2) = k(\boldsymbol{x}_1, \boldsymbol{x}_2)$. Therefore, we have $cov(y_1, y_2) = k(\boldsymbol{x}_1, \boldsymbol{x}_2) + \epsilon^2$. We then expand our objective function as

$$Var[(\hat{f}(\boldsymbol{x}) - F(\boldsymbol{x}))^2] = \kappa(\boldsymbol{x}, \boldsymbol{x}) + \boldsymbol{\lambda}(\boldsymbol{\kappa}(\boldsymbol{x}))^T(\boldsymbol{K} + \epsilon^2\boldsymbol{I})\boldsymbol{\lambda}(\boldsymbol{x}) - 2\boldsymbol{\lambda}(\boldsymbol{x})^T\boldsymbol{k}$$

where $\boldsymbol{K}$ is the covariance matrix with $[\boldsymbol{K}_n]_{ij} = k(\boldsymbol{x}_i, \boldsymbol{x}_j)$; $\boldsymbol{\kappa}(\boldsymbol{x})$ is the covariance vector between $\boldsymbol{x}$ and $\boldsymbol{x}_1, \boldsymbol{x}_2, ..., \boldsymbol{x}_n$, and $\boldsymbol{\lambda}(\boldsymbol{x})$ is the vector of $\lambda_1(\boldsymbol{x}), \lambda_2(\boldsymbol{x}), ..., \lambda_n(\boldsymbol{x})$.

In [Matheron, 1963], it assumes that $f(\boldsymbol{x})$ consists of a linear combination of finite low-degree functions:

$$f(\boldsymbol{x}) = \sum_{i=1}^{l} \beta_i f_i(\boldsymbol{x})$$

where the coefficient vector $\boldsymbol{\beta}$ is unknown. Then we could expand the unbiased constraint as:

$$\sum_{i}^{l}\sum_{j}^{n} \beta_i \lambda_j(\boldsymbol{x}) f_i(\boldsymbol{x}_j) = \sum_{i}^{l} \beta_i f_i(\boldsymbol{x})$$

Because the above equation should hold for any arbitrary $\boldsymbol{\beta}$, we obtain:

$$\sum_{j}^{l} \lambda_j(\boldsymbol{x}) f_i(\boldsymbol{x}_j) = f_i(\boldsymbol{x}) \text{ for all i}$$

We write it into the matrix form: $\boldsymbol{G}\boldsymbol{\lambda}(\boldsymbol{x}) = \boldsymbol{f}$, where $\boldsymbol{G}_{l\times n} = \begin{bmatrix} f_1(\boldsymbol{x}_1) & f_1(\boldsymbol{x}_2) & \dots \\ f_2(\boldsymbol{x}_1) & f_2(\boldsymbol{x}_2) & \dots \\ \dots & & \end{bmatrix}$, and $\boldsymbol{f}$ is the vector of $f_1(\boldsymbol{x}), f_2(\boldsymbol{x}), \dots$.

We let $\boldsymbol{\gamma}$ be the vector of the Lagrangian multiplier and write the Lagrangian formula:

$$L(\boldsymbol{\lambda}(\boldsymbol{x}), \gamma) = \kappa(\boldsymbol{x}, \boldsymbol{x}) + \boldsymbol{\lambda}(\boldsymbol{x})^T(\boldsymbol{K} + \epsilon^2 \boldsymbol{I})\boldsymbol{\lambda}(\boldsymbol{x}) - 2\boldsymbol{\lambda}(\boldsymbol{x})^T \boldsymbol{\kappa}(\boldsymbol{x}) + 2\gamma^T(\boldsymbol{G}\boldsymbol{\lambda}(\boldsymbol{x}) - \boldsymbol{f})$$

We take the partial derivatives with respect to $\lambda(\boldsymbol{x})$ and $\boldsymbol{\gamma}$ and let them equal to 0:

$$\frac{\partial L}{\partial \boldsymbol{\lambda}(\boldsymbol{x})} = 2(\boldsymbol{K} + \epsilon^2 \boldsymbol{I})\lambda(\boldsymbol{x}) - 2\boldsymbol{\kappa}(\boldsymbol{x}) + 2\boldsymbol{G}^T \boldsymbol{\gamma} = \boldsymbol{0}$$

$$\frac{\partial L}{\partial \gamma} = 2(\boldsymbol{G}\boldsymbol{\lambda}(\boldsymbol{x}) - \boldsymbol{f}) = \boldsymbol{0}$$

The above two equations form the general linear system for Kriging. In practice, we usually assume a prior estimator $f_0(\boldsymbol{x})$ for $f(\boldsymbol{x})$. We thus shift the mean away to consider a zero-mean case:

$$P(\boldsymbol{x}) = F(\boldsymbol{x}) - f(\boldsymbol{x}) \approx F(\boldsymbol{x}) - f_0(\boldsymbol{x})$$

We solve the linear system for $P(\boldsymbol{x})$. As the mean of $P(\boldsymbol{x})$ equal to 0, the matrix $\boldsymbol{G}$ is a zero matrix. It is straightforward to get: $\boldsymbol{\lambda}(\boldsymbol{x}) = (\boldsymbol{K} + \epsilon^2 \boldsymbol{I})^{-1}\boldsymbol{\kappa}(\boldsymbol{x})$. Remember the observed $y_i$ for $P(\boldsymbol{x})$ should be also shifted by $f_0(\boldsymbol{x_i})$. Therefore, we get the estimator for $E[P(\boldsymbol{x})]$:

$$\hat{p}(\boldsymbol{x}) = \boldsymbol{\kappa}(\boldsymbol{x})^T(\boldsymbol{K} + \epsilon^2 \boldsymbol{I})^{-1}(\boldsymbol{y} - \boldsymbol{f_0})$$

where $\boldsymbol{y}$ and $\boldsymbol{f_0}$ are the vector of $y_1, y_2, ..., y_n$ and $f_0(\boldsymbol{x}_1), f_0(\boldsymbol{x}_2), ..., f_0(\boldsymbol{x}_n)$, respectively. We finally add the prior back to the equation, and get our final estimator for $f(\boldsymbol{x})$:

$$\hat{f}(\boldsymbol{x}) = \boldsymbol{\kappa}(\boldsymbol{x})^T(\boldsymbol{K} + \epsilon^2 \boldsymbol{I})^{-1}(\boldsymbol{y} - \boldsymbol{f_0}) + f_0(\boldsymbol{x})$$

So far we have proved the regressor used in NCPD is biased and then derived our Kriging regressor. The unbiased property of Kriging regressor could let the estimated function capture the location of the optimal funnel more accurate. Moreover, if the variogram is known, the expected square error of the kriging regressor is no greater than that of the NPR estimator [Yakowitz and Szidarovszky, 1985]. Lastly, NCPD regressor suffers mostly from its high biasness at the boundary of the search space, because of the asymmetry of the kernel weights in such regions. This will cause the posterior may still remain a high probability value at the boundary, while the probability values of the outside regions are regarded as 0.

#### 1.3 Empirical Comparison

In order to be more rigorous, we put the empirical comparison between the NCPD and BAL here. We use the same testing functions as in the main text and the same method for posterior analysis to get the optimization and uncertainty quantification results. For the optimization, in Table S1, our BAL

|  | NCPD |  |  | BAL |  |  |
|---|---|---|---|---|---|---|
| Dimension | 2 | 6 | 12 | 2 | 6 | 12 |
| Rastrigin | 1.04(0.36) | 1.96(0.34) | 5.35(1.23) | 0.84(0.05) | 1.83(0.11) | 3.58(0.87) |
| Rosenbrock | 0.78(0.73) | 1.96(0.86) | 4.00(0.56) | 0.23(0.19) | 1.43(0.75) | 2.25(0.21) |
| Griewank | 2.73(0.51) | 4.45(1.05) | 6.01(2.24) | 1.65(1.42) | 4.35(1.01) | 4.11(1.28) |
| Ackley | 0.33(0.31) | 0.91(0.25) | 2.63(0.31) | 0.26(0.14) | 0.69(0.23) | 1.69(0.11) |

Table S1: The means and the standard deviations (in the parenthesis) of $||\hat{\boldsymbol{x}} - \boldsymbol{x}^*||_2$ for four test functions on three dimensions.

|  |  | NCPD |  |  | BAL |  |  |
|---|---|---|---|---|---|---|---|
| Dim | | d | *por* | $T$ | d | *por* | $T$ |
| 2 | Rastrigin | 1.84(1.03) | 0.78 | 0.78(0.23) | 1.37(1.18) | 0.91 | 0.54(0.13) |
|  | Rosenbrock | 1.31(0.32) | 0.89 | 0.57(0.15) | 0.63(0.14) | 0.98 | 0.55(0.06) |
|  | Griewank | 3.91(1.88) | 0.80 | 0.65(0.22) | 2.85(1.45) | 0.86 | 0.62(0.20) |
|  | Ackley | 0.51(0.17) | 0.99 | 0.53(0.21) | 0.41(0.20) | 0.99 | 0.43(0.02) |
| 6 | Rastrigin | 2.68(1.54) | 0.72 | 0.33(0.12) | 2.25(1.02) | 0.91 | 0.22(0.11) |
|  | Rosenbrock | 3.04(1.98) | 0.90 | 0.51(0.16) | 2.16(1.43) | 0.92 | 0.50(0.25) |
|  | Griewank | 5.21(0.87) | 0.68 | 0.14(0.03) | 4.53(0.78) | 0.76 | 0.04(0.02) |
|  | Ackley | 1.13(0.22) | 0.89 | 0.33(0.30) | 0.77(0.34) | 0.96 | 0.20(0.08) |
| 12 | Rastrigin | 6.53(1.98) | 0.70 | 0.23(0.11) | 3.62(0.36) | 0.73 | 0.01(0.01) |
|  | Rosenbrock | 4.50(1.30) | 0.83 | 0.12(0.10) | 2.22(1.18) | 0.87 | 0.03(0.01) |
|  | Griewank | 7.05(2.23) | 0.67 | 0.11(0.03) | 4.25(1.01) | 0.63 | 0.05(0.03) |
|  | Ackley | 4.89(1.64) | 0.95 | 0.88(0.69) | 2.89(0.49) | 0.93 | 0.66(0.23) |

Table S2: Uncertainty Quantification for the test functions.

outperforms NCPD for every test function with every tested dimensionality. For UQ, in Table S2, the tightness of BAL is much lower then NCPD. At the same time, in majority of the cases, the portion of BAL is closer to 0.9. This means we have more tighter confidence interval but the accuracy of the interval is increasing at the same time. This is because our $\rho$ captured the global uncertainty in the system and is also dimensional independent, and we use a mini-batch sampling which could not only explore more on the search space, but also make the data more i.i.d. which will benefit the convergence of the regressor.

##### 1.4 Summary

We compare the posteriors of NCPD and our BAL from both theoretical and empirical prospectives. We claim three drawbacks of the $\rho$ in NCPD and how our BAL overcomes it. We shows the regressor we used is the best linear unbiased regressor. The empirical results show our method outperforms theirs in both optimization and uncertainty quantification.

#### 2 Methods

##### 2.1 Fast RMSD calculation

We describe how we calculate the interface RMSD (iRMSD) between two sample structures in $O(1)$ instead of $O(N)$, where $N$ is the number of interfacial atoms. We will also show this method is generalizable for the calculation of any kind of RMSD and the case that the rigid-body motion and the flexibility are considered separately.

Assume $\boldsymbol{C_1^{int}}$ and $\boldsymbol{C_2^{int}}$ are the vectors of the coordinates of interfacial atoms of two docking complexes, respectively:

$$
\begin{aligned}
\boldsymbol{C_1^{int}} &= \boldsymbol{C_0^{int}} + \sum_{i=1}^{d} r_i \boldsymbol{\mu_i^{int}} \\
\boldsymbol{C_2^{int}} &= \boldsymbol{C_0^{int}} + \sum_{i=1}^{d} r'_i \boldsymbol{\mu_i^{int}}
\end{aligned} \tag{1}
$$

where d is the dimension of search space and $\boldsymbol{C_0}$ is the vector of the coordinates of interfacial atoms of the starting structure. $r_i$ and $r'_i$ are the coefficients before normal mode i of two docking complexes, respectively, and $\boldsymbol{\mu_i}$ is the interface-specific subvector of complex normal mode i.

The iRMSD between these two structures is:

$$
\begin{aligned}
iRMSD = \sqrt{\frac{||\boldsymbol{C_1^{int}} - \boldsymbol{C_2^{int}}||^2}{N}} &= \sqrt{\frac{||\sum_{i=1}^{d}(r_i - r'_i)\boldsymbol{\mu_i^{int}}||^2}{N}} \\
&= \sqrt{\frac{\sum_{i=1}^{d}\sum_{j=1}^{d}(r_i - r'_i)(r_j - r'_j)\boldsymbol{\mu_i^{int}} \cdot \boldsymbol{\mu_j^{int}}}{N}}
\end{aligned} \tag{2}
$$

Let $\boldsymbol{U}^{int}$ be the matrix of $[\boldsymbol{U}^{int}]_{ij} = \boldsymbol{\mu_i^{int}} \cdot \boldsymbol{\mu_j^{int}}$, and $\boldsymbol{r}$ be the row vector of $[\boldsymbol{r}]_i = (r_i - r'_i)$, then the iRMSD could be rewritten as:

$$
iRMSD = \sqrt{\frac{\boldsymbol{r}^T \boldsymbol{U} \boldsymbol{r}}{N}} \tag{3}
$$

Because $\boldsymbol{U}$ is a $d \times d$ matrix, we could calculate the iRMSD in $O(d^2) = O(1)$.

It is straightforward to extend this equation for other kinds of RMSD. For the case that rigid-body motion is separated from the flexibility, we suppose the rotation matrix for one atom is:

$$
\boldsymbol{w} = \begin{bmatrix} w_1 & w_2 & w_3 \\ w_4 & w_5 & w_6 \\ w_7 & w_8 & w_9 \end{bmatrix} \tag{4}
$$

We extend it to all-atom case:

$$
\boldsymbol{W} = [\underbrace{\boldsymbol{w}, \boldsymbol{w}, ..., \boldsymbol{w}}_{\text{N}}] \tag{5}
$$

More generally, We consider $\boldsymbol{C_1}$ and $\boldsymbol{C_2}$ as the vector of atoms for two sample structures:

$$
\begin{aligned}
\boldsymbol{C_1} &= \boldsymbol{C_{center}} + \sum_{i=1}^{3} t_i \boldsymbol{T_i} + \boldsymbol{W}(\boldsymbol{C_0} - \boldsymbol{C_{center}}) + \sum_{i=1}^{d} r_i \boldsymbol{\mu_i} \\
\boldsymbol{C_2} &= \boldsymbol{C_{center}} + \sum_{i=1}^{3} t'_i \boldsymbol{T_i} + \boldsymbol{W'}(\boldsymbol{C_0} - \boldsymbol{C_{center}}) + \sum_{i=1}^{d} r'_i \boldsymbol{\mu_i}
\end{aligned} \tag{6}
$$

where $\boldsymbol{C_{center}}$ is the rotation center of the starting structure, and $t_i$s are the translational coefficients, and $\boldsymbol{T_1}$ is the $\mathbf{1}$ vector for $x$ dimension:

$$
\boldsymbol{T_1} = [\underbrace{1\,0\,0\,1\,0\,0 ...}_{\text{3N}}]^T
$$

$\boldsymbol{T_2}$ and $\boldsymbol{T_3}$ are in the same manner for $y$ and $z$ dimensions.

It is easy to rewrite the rotation part in the way as:

$$
\begin{aligned}
\boldsymbol{C_1} &= \boldsymbol{C_{center}} + \sum_{i=1}^{3} t_i \boldsymbol{T_i} + \sum_{i=1}^{i=9} w_i \boldsymbol{c_i} + \sum_{i=1}^{d} r_i \boldsymbol{\mu_i} \\
\boldsymbol{C_2} &= \boldsymbol{C_{center}} + \sum_{i=1}^{3} t'_i \boldsymbol{T_i} + \sum_{i=1}^{i=9} w'_i \boldsymbol{c_i} + \sum_{i=1}^{d} r'_i \boldsymbol{\mu_i}
\end{aligned} \tag{7}
$$

where if we let $\boldsymbol{C_0} - \boldsymbol{C_{center}} = [C_{1x}, C_{1y}, C_{1z}, C_{2x} ... C_{Nx}, C_{Ny}, C_{Nz}]$, then

$$\begin{aligned} \boldsymbol{c_1} &= [C_{1x}, 0, 0, ..., C_{Nx}, 0, 0], \;\; \boldsymbol{c_2} = [C_{1y}, 0, 0, ..., C_{Ny}, 0, 0], \;\; \boldsymbol{c_3} = [C_{1z}, 0, 0, ..., C_{Nz}, 0, 0], \\ \boldsymbol{c_4} &= [0, C_{1x}, 0, ..., 0, C_{Nx}, 0], \;\; \boldsymbol{c_5} = [0, C_{1y}, 0, ..., 0, C_{Ny}, 0], \;\; \boldsymbol{c_6} = [0, C_{1z}, 0, ..., 0, C_{Nz}, 0], \\ \boldsymbol{c_7} &= [0, 0, C_{1x}, ..., 0, 0, C_{Nx}], \;\; \boldsymbol{c_8} = [0, 0, C_{1y}, ..., 0, 0, C_{Ny}], \;\; \boldsymbol{c_9} = [0, 0, C_{1z}, ..., 0, 0, C_{Nz}], \end{aligned} \tag{8}$$

Similar to the previous discussion, the RMSD between $\boldsymbol{C_1}$ and $\boldsymbol{C_2}$ could be calculated in $O((3+9+d) \times (3+9+d)) = O(d^2) = O(1)$.

#### 2.2 Distribution of the ratio between predicted and actual extents of conformational changes for the receptor

We describe the distribution in this section for range reduction in the reduced conformational space. In our previous work [Chen et al., 2017], we predicted the extent of conformational changes of receptor for an encounter complex as $\widehat{\text{RMSD}}_{\text{R}}$. We then calculate $\text{RMSD}_{\text{R}}/\widehat{\text{RMSD}}_{\text{R}}$ for the 500 models in the training set and fit it into a truncated Gaussian distribution, which is shown in Fig. S2.

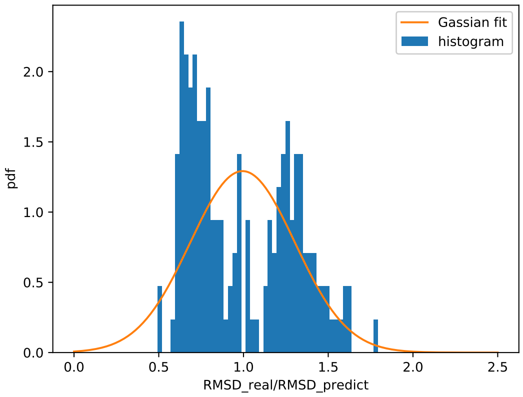


Figure S2: The histogram of the ratio. The fit Gaussian distribution has mean and standard deviation equal to 0.99 and 0.31, respectively.

This distribution, multiplying $\widehat{\text{RMSD}}_{\text{R}}$, will be later used as the prior distribution of $\tau_{\text{R}}$ for sampling initial 30 structures.

#### 2.3 Feasibility of the search space for sampling

We consider the feasibility of the search space here. For samples generated from the updated posterior, they have only one constraint which is

$$\sqrt{\frac{1}{N_{\text{L}}}} \left\| \sum_{j \in \mathcal{B}} r_j \frac{s}{\sqrt{\lambda_j}} \cdot \boldsymbol{\mu}_j^{\text{L}} \right\| \leqslant \overline{\Delta}_{\text{L}} \tag{9}$$

If we replace $r_j \cdot s$ with our parameterization $x_j$, it is obvious to see the search space is within a ellipsoid ball.

For samples generated from the prior, besides the aforementioned constraint, its another inequality constraint is given by:

$$\sqrt{\frac{1}{N_{\text{R}}}} \left\| \sum_{j \in \mathcal{B}} r_j \frac{s}{\sqrt{\lambda_j}} \cdot \boldsymbol{\mu}_j^{\text{R}} \right\| \leqslant 2.5 \cdot \widehat{\text{RMSD}}_{\text{R}} \tag{10}$$

where 2.5 is from the upper bound of our truncated normal distribution (see Fig. S2). Therefore, the search space is the intersection of the aforementioned two inequality constraints. It is obvious that the origin is a feasible point. The above constraints hereby have feasible solutions. Lastly, just for the consideration of running time, during the prior sampling, the $\widehat{\text{RMSD}}_{\text{R}}$ will be reduced until the acceptance rate of the reject sampling is found to be no less than a threshold (0.001%).

#### 2.4 Putative interface

We briefly introduce how to get the putative interface for each starting structure (the starting representative of each region). Specifically, each starting complex structure is perturbed based on the prior distribution of $\boldsymbol{x}$ for 50 times. The interface residues are chosen for each perturbed complex with intermolecular atomic distance cutoff equal to 10Å. And then the union of 50 interface sets is regarded as the putative interface for this model.

#### 2.5 List of protein complexes used in the study

| Difficulty | Index | PDB ID | Index | PDB ID |
|---|---|---|---|---|
| Rigid | 1 | 1N8O | 19 | 1EAW |
| | 2 | 7CEI | 20 | 2JEL |
| | 3 | 1DFJ | 21 | 1ML0 |
| | 4 | 1AVX | 22 | 1BJ1 |
| | 5 | 1AY7 | 23 | 1KXQ |
| | 6 | 1BVN | 24 | 1EWY |
| | 7 | 1IQD | 25 | 1KAC |
| | 8 | 1CGI | 26 | 1OPH |
| | 9 | 1MAH | 27 | 2AJF |
| | 10 | 1EZU | 28 | 1E6J |
| | 11 | 1JPS | 29 | 2HLE |
| | 12 | 1PPE | 30 | 1WEJ |
| | 13 | 1R0R | 31 | 1A2K |
| | 14 | 1T6B | 32 | 1RLB |
| | 15 | 2FD6 | 33 | 1GLA |
| | 16 | 2I25 | 34 | 1E6E |
| | 17 | 2B42 | 35 | 1J2J |
| | 18 | 1BUH | | |
| Medium | 1 | 1XQS | 5 | 1IJK |
| | 2 | 1BGX | 6 | 2HRK |
| | 3 | 1KKL | 7 | 1GP2 |
| | 4 | 1M10 | 8 | 1GRN |
| Flexible | 1 | 1IBR | 5 | 1H1V |
| | 2 | 1BKD | 6 | 1DE4 |
| | 3 | 1Y64 | 7 | 1ATN |
| | 4 | 2C0L | | |

Table S3: Protein complexes in the training set for scoring function and the extents of conformational changes. $K_d$ values are known.

| Difficulty | Index | PDB ID | Index | PDB ID |
|---|---|---|---|---|
| Rigid | 1 | 1AHW | 44 | 1PVH |
| | $2^\dagger$ | 1AK4 | $45^\dagger$ | 1QA9 |
| | $3^\dagger$ | 1AKJ | 46 | 1QFW |
| | $4^\dagger$ | 1AZS | 47 | 1RV6 |
| | $5^\dagger$ | 1B6C | $48^\dagger$ | 1S1Q |
| | 6 | 1BVK | $49^\dagger$ | 1SBB |
| | 7 | 1CLV | 50 | 1TMQ |
| | 8 | 1D6R | 51 | 1UDI |
| | 9 | 1DQJ | 52 | 1US7 |
| | $10^\dagger$ | 1E96 | 53 | 1VFB |
| | $11^\dagger$ | 1EFN | 54 | 1WDW |
| | 12 | 1F34 | 55 | 1XD3 |
| | 13 | 1F51 | 56 | 1XU1 |
| | $14^\dagger$ | 1FC2 | 57 | 1YVB |
| | 15 | 1FCC | $58^\dagger$ | 1Z0K |

| | | | | |
|---|---|---|---|---|
| | 16 | 1FFW | 59 | 1Z5Y |
| | 17 | 1FLE | 60 | 1ZHH |
| | 18 | 1FQJ | 61 | 1ZHI |
| | 19 | 1FSK | 62 | 2A5T |
| | 20† | 1GCQ | 63 | 2A9K |
| | 21† | 1GHQ | 64 | 2ABZ |
| | 22 | 1GL1 | 65 | 2B4J |
| | 23† | 1GPW | 66† | 2BTF |
| | 24 | 1GXD | 67 | 2FJU |
| | 25 | 1H9D | 68 | 2G77 |
| | 26 | 1HCF | 69† | 2HQS |
| | 27 | 1HE1 | 70 | 2J0T |
| | 28 | 1HIA | 71† | 2MTA |
| | 29† | 1I4D | 72 | 2O8V |
| | 30 | 1I9R | 73 | 2OOB |
| | 31 | 1JTG | 74 | 2OOR |
| | 32 | 1JWH | 75 | 2OUL |
| | 33 | 1K4C | 76† | 2PCC |
| | 34 | 1K74 | 77 | 2SIC |
| | 35 | 1KLU | 78 | 2SNI |
| | 36 | 1KTZ | 79 | 2UUY |
| | 37 | 1KXP | 80 | 2VDB |
| | 38 | 1MLC | 81 | 2VIS |
| | 39 | 1NCA | 82 | 3BP8 |
| | 40 | 1NSN | 83 | 3D5S |
| | 41 | 1OC0 | 84 | 3SGQ |
| | 42 | 1OFU | 85 | 9QFW |
| | 43 | 1OYV | 86 | BOYV |
| Medium | 1 | 1ACB | 12 | 1SYX |
| | 2 | 1HE8 | 13† | 1WQ1 |
| | 3† | 1I2M | 14 | 2AYO |
| | 4† | 1IB1 | 15 | 2CFH |
| | 5 | 1JIW | 16 | 2H7V |
| | 6 | 1K5D | 17 | 2J7P |
| | 7 | 1LFD | 18 | 2NZ8 |
| | 8 | 1MQ8 | 19 | 2OZA |
| | 9 | 1N2C | 20 | 2Z0E |
| | 10† | 1NW9 | 21 | 3CPH |
| | 11 | 1R6Q | 22 | 4CPA |
| Flexible | 1 | 1E4K | 10 | 1PXV |
| | 2 | 1EER | 11† | 1R8S |
| | 3 | 1F6M | 12 | 1ZLI |
| | 4† | 1FAK | 13 | 1ZM4 |
| | 5 | 1FQ1 | 14 | 2HMI |
| | 6 | 1IRA | 15 | 2I9B |
| | 7 | 1JK9 | 16 | 2IDO |
| | 8 | 1JMO | 17 | 2O3B |
| | 9 | 1JZD | 18† | 2OT3 |

Table S4: Protein complexes in the test set for both scoring function and protein docking. †: Protein complexes with known $K_d$ values.

#### 2.6 Energy Model Training

##### 2.6.1 Training set

The whole dataset contains 10 encounter complex structures for each of the aforementioned 176 protein pairs (Sec. 2.5 Tables S3 and S4) in the Protein Docking Benchmark Set 4.0 [Hwang et al., 2010]. They were generated by ZDOCK as the top 10 cluster centers and kindly provided by the Weng group. 50 of

| Index | PDB ID | $\text{iRMSD}_\text{C}$(Å) |
|---|---|---|
| 1 | 2REX | 1.107 |
| 2 | 2WPT | 1.609 |
| 3 | 3BX1 | 1.100 |
| 4 | 3FM8 | 1.818 |
| 5 | 3Q87 | 3.739 |
| 6 | 4G9S | 3.739 |
| 7 | 4JW2 | 2.929 |
| 8 | 4JW3 | 1.926 |
| 9 | 4OJK | 1.832 |
| 10 | 4QKO | 0.950 |
| 11 | 4QT8 | 1.451 |
| 12 | 4UEM | 8.102 |
| 13 | 4UF5 | 16.890 |
| 14 | 4UHP | 1.891 |
| 15 | 4XL5 | 3.223 |

Table S5: Protein complexes in CAPRI test set for protein docking. $\text{iRMSD}_\text{C}$ here is the interface RMSD after superimposing unbound receptor and ligand to the bound receptor and ligand separately. Higher $\text{iRMSD}_\text{C}$ suggests more conformational changes upon protein-protein interactions and more challenges to protein docking. $K_d$ values are predicted from sequence alone.

176 protein pairs (See Table S3) have been chosen as the training set for training the scoring function, including 35(70%) rigid cases, 8(16%) medium cases and 7(14%) flexible cases. The $K_d$ values of these 50 targets are provided by the Binding Affinity Benchmark Set [Kastritis and Bonvin, 2010]. Within all 500 models, 64 models are near-native ones (iRMSD$\leq$ 4 Å). In order to balance the ratio of near-native and non near-native models in the training set to improve the performance for the near-native part, we use a strategy called oversampling to balance the training data. Specifically, each of 436 non-near native models are perturbed for 15 times and each of 64 near-native models are perturbed for 101 times. In all, we have 13,004 examples made up of 6,540 non-near native examples and 6,464 near-native examples in the training set. Notice here, the way to perturb each model is consistent with that in our docking. The rationale is that the sample distribution in the training set needs to be consistent with the sample distribution in the docking process. Otherwise, others need to do some transformed learning.
All the energy feature values were standardized before training. Random forest and Ridge regression with linear and nonlinear radial basis function (RBF) kernel were performed with 4-fold cross validation over the training set to determine hyperparameters and model parameters. Specifically, the hyperparameters were determined by searching on discrete grids for the optimal values that minimize mean squared errors (MSE) averaged over all 4 folds.

###### 2.6.2 Hyperparameter Tuning

The internal hyperparameters of each machine learning model can be tuned through the cross-validation. One technical issue here is that, for $\alpha$ and $q$ which are made up of the label, it is not quite trivial to be tuned through cross-validation because the labels are different for different sets of $\alpha$ and $q$, so that the traditional scores like Mean Square Error(MSE) or Pearson's $r$ are not comparable across different sets of $\alpha$ and $q$. We need to find a common assessment metric which is independent of $\alpha$ and $q$. To reach this goal, we used the mean of Spearman correlation over the 50 training protein pairs between the labels and the iRMSDs. The reason to use Spearman correlation is because, first, the monotonicity or the ranking ability is our goal for training this scoring function. Secondly, Spearman correlation could be comparable across different sets of $\alpha$ and $q$ within each protein pair. The mean of the Spearman correlation could be hereby used as a universal metric for optimizing $\alpha$ and $q$ in the cross-validation. The optimal $\alpha$ and $q$ after training are shown in Table S10.

##### 2.7 PSO and BAL for Test Functions

For particle swarm optimization (PSO), we have used the standard version with inertia correction [Xu et al., 2007, Clerc and Kennedy, 2002, Bansal et al., 2011]. In order to be fair, the swarm size and the number of iterations are the same for both algorithms, which are shown in Table S6. Either algorithm is run for 100 times for the statistical significance. The search regions are shown in Table S7.

| Dimension | 2 | 6 | 12 |
|---|---|---|---|
| Swarm-size | 10 | 15 | 20 |
| Iterations | 50 | 100 | 150 |
| Total samples | 500 | 1500 | 3000 |

Table S6: Parameters for BAL and PSO algorithms

| | |
|---|---|
| Rastrigin | $-8 \leqslant x_i \leqslant 8,\ 1 \leqslant i \leqslant d$ |
| Rosenbrock | $-8 \leqslant x_i \leqslant 8,\ 1 \leqslant i \leqslant d$ |
| Griewank | $-8 \leqslant x_i \leqslant 8,\ 1 \leqslant i \leqslant d$ |
| Ackley | $-20 \leqslant x_i \leqslant 20,\ 1 \leqslant i \leqslant d$ |

Table S7: Search regions for test function.

### 3 Results

#### 3.1 Performances on Test Functions

| | PSO | | | BAL | | |
|---|---|---|---|---|---|---|
| Dimension $d$ | 2 | 6 | 12 | 2 | 6 | 12 |
| Rastrigin | 0.99 (0.56) | 2.06 (0.15) | 4.06 (1.01) | 0.84 (0.05) | 1.83 (0.11) | 3.58 (0.87) |
| Rosenbrock | 0.91 (0.64) | 2.08 (0.86) | 3.12 (0.33) | 0.23 (0.19) | 1.43 (0.75) | 2.25 (0.21) |
| Griewank | 3.63 (0.33) | 5.42 (1.01) | 5.51 (2.29) | 1.65 (1.42) | 4.35 (1.01) | 4.11 (1.28) |
| Ackley | 0.37 (0.21) | 0.88 (0.26) | 2.65 (0.38) | 0.26 (0.14) | 0.69 (0.23) | 1.69 (0.11) |

Table S8: Optimization performances of PSO and BAL over four non-convex test functions in various dimensions based on means (and standard deviations in parentheses) of $||\hat{\boldsymbol{x}} - \boldsymbol{x}^*||$, the distance between the predicted and the actual global optima.

| Dimension $d$ | Function | $r_{90}$ | $\eta$ | $\hat{P}$ |
|---|---|---|---|---|
| 2 | Rastrigin | 1.37 (1.18) | 0.54 (0.13) | 0.91 |
| | Rosenbrock | 0.40 (0.24) | 0.55 (0.06) | 0.98 |
| | Griewank | 2.85 (1.45) | 0.62 (0.20) | 0.86 |
| | Ackley | 0.41 (0.20) | 0.43 (0.02) | 0.99 |
| 6 | Rastrigin | 2.25 (1.02) | 0.22 (0.11) | 0.91 |
| | Rosenbrock | 2.16 (1.43) | 0.50 (0.25) | 0.92 |
| | Griewank | 4.53 (0.78) | 0.04 (0.02) | 0.76 |
| | Ackley | 0.77 (0.34) | 0.20 (0.08) | 0.96 |
| 12 | Rastrigin | 3.62 (0.36) | 0.01 (0.01) | 0.73 |
| | Rosenbrock | 2.22 (1.18) | 0.03 (0.01) | 0.87 |
| | Griewank | 4.25 (1.01) | 0.05 (0.03) | 0.63 |
| | Ackley | 2.89 (0.49) | 0.66 (0.23) | 0.93 |

Table S9: Uncertainty quantification performances of BAL over test functions based on $r_{90}$, the estimated upper bound of $||\hat{\boldsymbol{x}} - \boldsymbol{x}^*||$ at a 90% confidence level; $\eta$, the relative error in $r_{90}$; and $\hat{P}$, the portion of confidence intervals from 100 runs encompassing the global optima. For $r_{90}$ and $\eta$, means (and standard deviations in parentheses) are reported.

#### 3.2 Energy Model for Protein Docking

| Models | $\alpha$ | $q$ |
|---|---|---|
| Ridge | 6.70 | 0.75 |
| Ridge with RBF | 12.0 | 0.50 |
| Random Forest | 8.72 | 0.50 |

Table S10: Optimal $\alpha$ and $q$ for different machine learning models after training.

| | Ridge | Ridge with RBF | Random Forest |
|---|---|---|---|
| Training | 8.32(0.25) | 5.45(0.67) | 2.45(0.79) |
| Test | 12.34(0.26) | 10.34(0.67) | 4.78(0.75) |

Table S11: Performance on training and test **native** sets based on RMSE (and Pearson correlation in parentheses) between predicted $y(\boldsymbol{x})$ and real $y(\boldsymbol{x})$. The unit of RMSE is Kcal/mol.

#### 3.3 Protein docking

##### 3.3.1 Comparison between PSO and BAL on energy scores ($y$)

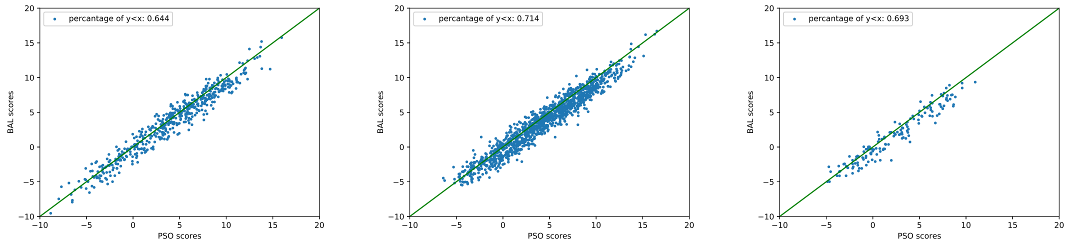


(a) (b) (c)

Figure S3: Head to head comparison between BAL and PSO predictions $\hat{\mathbf{x}}$ in (random forest) energy scores $y(\hat{\mathbf{x}}_i)$ for training (a), test (b) and CAPRI (c) sets.

##### 3.3.2 Comparison between PSO and BAL on solution quality (iRMSD)

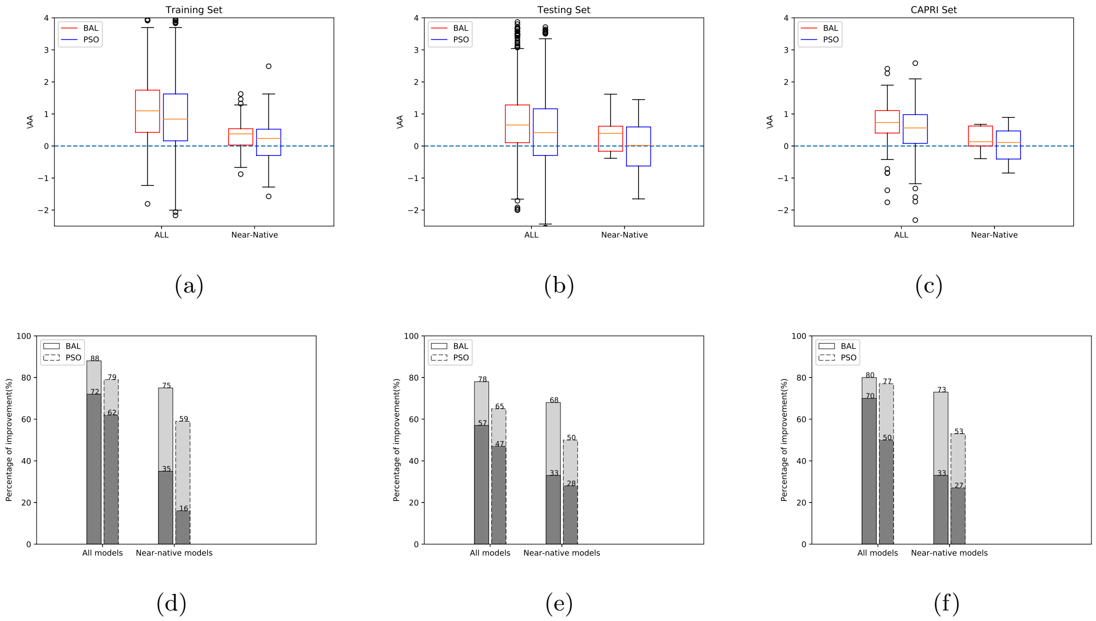


Figure S4: Box plots for the improvement in RMSD after PSO and BAL refinements for the training (a), test (b) and CAPRI (c) sets. Also reported are the percentages of BAL (solid bars) and PSO (dashed bars) refinement results with iRMSD improvement or with significant iRMSD improvement over 0.5Å(the darker portions) for the training (d), test (e) and CAPRI (f) sets.

##### 3.3.3 Percentage improvement

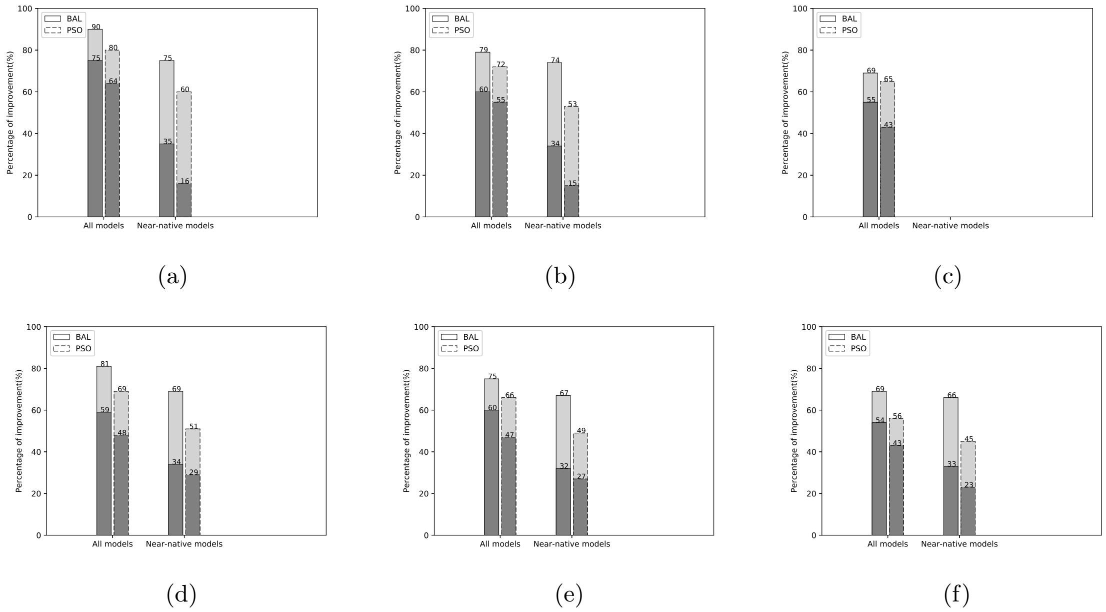


Figure S5: The relative percentage improvement against the starting structure of BAL and PSO for (a) Rigid cases in the training set. (b) Medium cases in the training set. (c)Flexible cases in the training set. (d) Rigid cases in the testing set. (e) Medium cases in the testing set. (f) Flexible cases in the testing set.

#### 3.4 Sampled Energy Landscapes

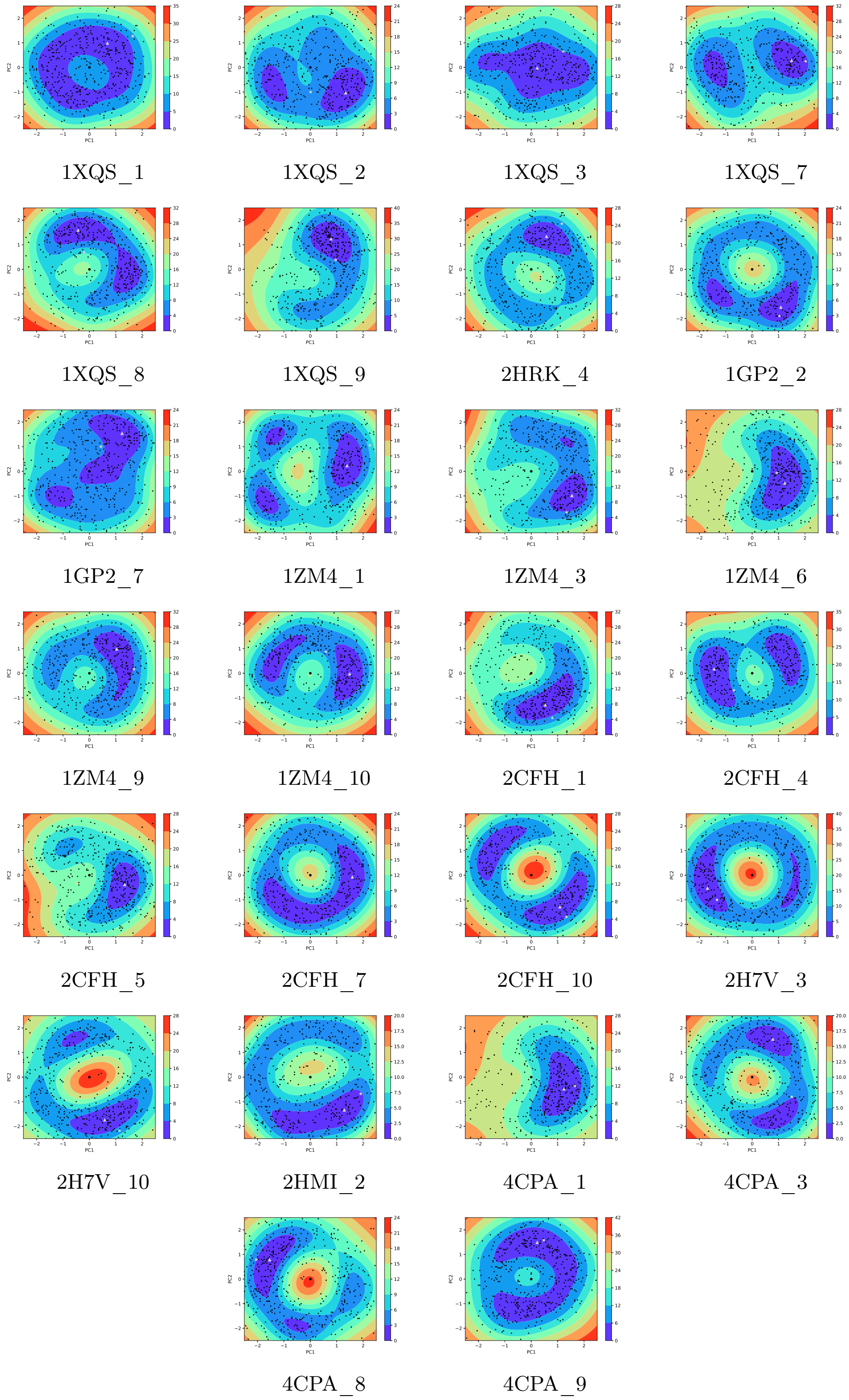


Figure S6: The estimated energy landscapes along the first and the second principle components (PC) for near-native and medium or difficult models in the **benchmark** set (training set and test set). All the black dots are the samples. The grey triangle is the estimated end structure and the grey star is the true native structure. The starting structure is a thicker black dot at the origin. All the energy values are re-centered to let the lowest energy value equal to 0 within each model.

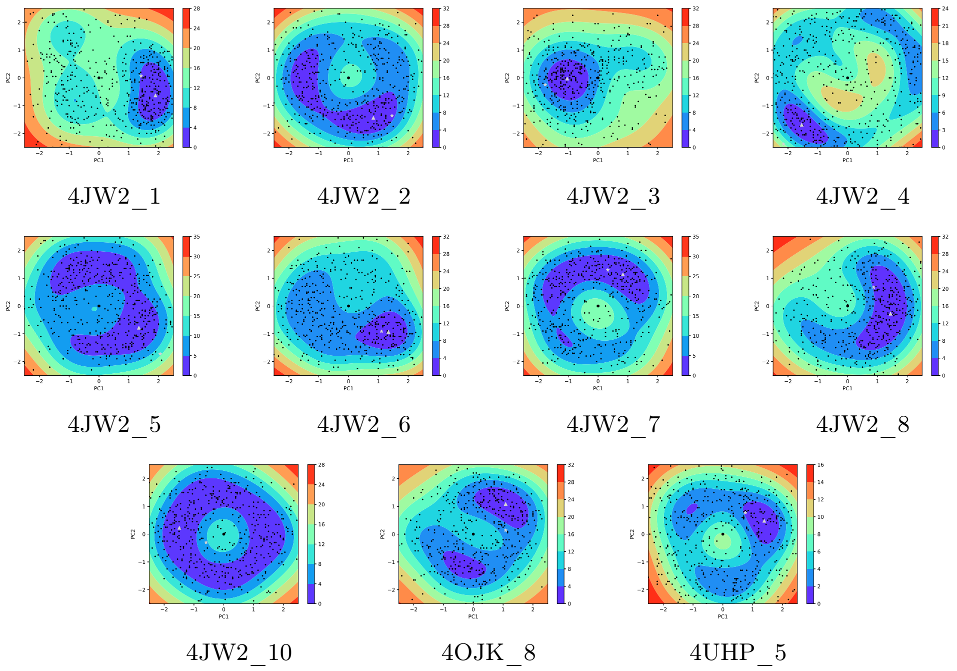


Figure S7: The estimated energy landscapes along the first and the second principle components (PC) for near-native and medium or difficult models in the **CAPRI** set. All the black dots are the samples. The grey triangle is the estimated end structure and the grey star is the true native structure. The starting structure is a thicker black dot at the origin. All the energy values are re-centered to let the lowest energy value equal to 0 within each model.

##### 3.5 List of parameter values used in this paper

| Parameters | $\rho_0$ | $l_0$ | $\epsilon$ |
|---|---|---|---|
| Values | 1.0 | 2.0 | 2.1 |

Table S12: Parameters used for BAL Implementation

##### 3.6 Running time for optimization

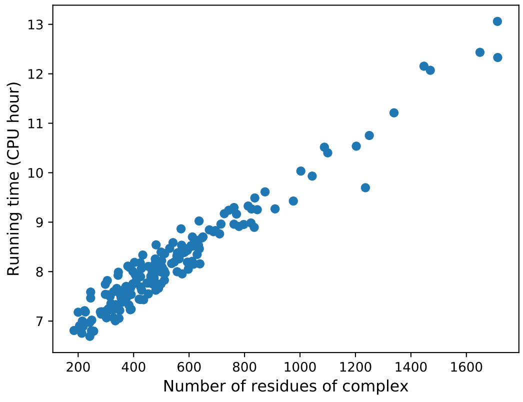


Figure S8: Running time (core hours) for BAL optimization for each 630-sample refinement on Intel Xeon 2.5GHz E5-2670. Post-optimization UQ takes 0.5 to 1 additional core hour each.

#### 4 Videos

We also attach with the supporting information three videos to show actual BAL optimization trajectories for protein docking and one video to showcase the slowest complex normal mode blending flexible-body motions of both the receptor and the ligand and rigid-body motions of the ligand.

Jean-Paul Chilès and Pierre Delfiner. *Geostatistics: Modeling Spatial Uncertainty, 2nd Edition.* 2012.

Maurice Clerc and James Kennedy. The particle swarm-explosion, stability, and convergence in a multidimensional complex space. *IEEE transactions on Evolutionary Computation*, 6(1):58–73, 2002.

Howook Hwang, Thom Vreven, JoÃńl Janin, and Zhiping Weng. Protein-Protein Docking Benchmark Version 4.0. *Proteins*, 78(15):3111–3114, November 2010. ISSN 0887-3585. doi: 10.1002/prot.22830. URL https://www.ncbi.nlm.nih.gov/pmc/articles/PMC2958056/.

Panagiotis L. Kastritis and Alexandre M. J. J. Bonvin. Are Scoring Functions in ProteinProtein Docking Ready To Predict Interactomes? Clues from a Novel Binding Affinity Benchmark. *Journal of Proteome Research*, 9(5):2216–2225, May 2010. ISSN 1535-3893. doi: 10.1021/pr9009854. URL https://doi.org/10.1021/pr9009854.

Georges Matheron. Principles of geostatistics. *Economic geology*, 58(8):1246–1266, 1963.

Pedro Ortega, Jordi Grau-Moya, Tim Genewein, David Balduzzi, and Daniel Braun. A nonparametric conjugate prior distribution for the maximizing argument of a noisy function. In *Advances in Neural Information Processing Systems*, pages 3005–3013. IEEE, 2012.

Rui Xu, Donald Wunsch II, and Ronald Frank. Inference of genetic regulatory networks with recurrent neural network models using particle swarm optimization. *IEEE/ACM Transactions on Computational Biology and Bioinformatics*, 4(4):681–692, 2007.

SJ Yakowitz and F Szidarovszky. A comparison of kriging with nonparametric regression methods. *Journal of Multivariate Analysis*, 16(1):21–53, 1985.
